# Characterization of human XPD helicase activity with Single Molecule Magnetic Tweezers

**DOI:** 10.1101/2023.02.07.527508

**Authors:** Chunfeng Mao, Maria Mills

## Abstract

XPD helicase is a DNA unwinding enzyme involved in multiple cellular processes. As part of TFIIH, XPD opens a repair bubble in DNA for access by proteins in the nucleotide excision repair pathway. XPD uses the energy from ATP hydrolysis to translocate in the 5’ to 3’ direction on one strand of duplex DNA, displacing the opposite strand in the process. We used magnetic tweezers assays to measure the double-stranded DNA (dsDNA) unwinding and single-stranded DNA (ssDNA) translocation activities of human XPD by itself. In our experimental setup, hXPD exhibits low unwinding processivity of ~14 bp and slow overall unwinding rate of ~0.3 bp/s. Individual unwinding and translocation events were composed of fast and slow runs and pauses. Analysis of these events gave similar mean run sizes and rates for unwinding and translocation, suggesting that unwinding is a reflection of translocation. The analysis also revealed that hXPD spent similar time stalling and unwinding. hXPD translocated on ssDNA at a similar overall rate as that of unwinding, pointing to an active helicase. However, we observed modest effects of DNA sequence on stalling and unwinding initiation position. Considering the slow unwinding rate, high probability of base pair separation at the ssDNA/dsDNA fork, and the observed DNA sequence dependences, we propose that hXPD is most likely a partially active helicase. Our results provide detailed information on the basal activity of hXPD which enhances our mechanistic understanding of hXPD activity.

**SIGNIFICANCE:** Human XPD helicase is a major component of the general transcription factor TFIIH. TFIIH is essential in both transcription and nucleotide excision repair. Mutations in hXPD are associated with cancers and autosomal recessive disorders. Here we directly measured the dsDNA unwinding and ssDNA translocation of human XPD helicase by itself. Our measurements provide detailed information on the basal activity of human XPD, which enhance our mechanistic understanding of the activity of XPD in the cell, provide a basis for better understanding of the clinical phenotypes, and aid in drug design targeting hXPD related diseases.

## INTRODUCTION

Human XPD (hXPD) is an SF2 family helicase with diverse roles in the cell, including DNA repair, transcription, and cell cycle regulation (1–3). XPD is composed of two RecA-like domains, HD1 and HD2, an Arch domain, and a fourth domain containing a FeS cluster (Figure 1A, structure adapted from PDB 6RO4 (4)). XPD unwinds DNA by translocating on ssDNA from 5’ to 3’ using the energy of ATP hydrolysis (5). The most studied role of XPD is within the general transcription factor TFIIH complex in the nucleus. TFIIH is a large multidomain complex which plays a critical role in both transcription initiation and nucleotide excision repair (NER) pathways. TFIIH opens DNA for access by RNA polymerase II and NER enzymes. The helicase activity of XPD is required for the NER pathway, while its role in transcription is structural (6–8).

**Figure 1.**
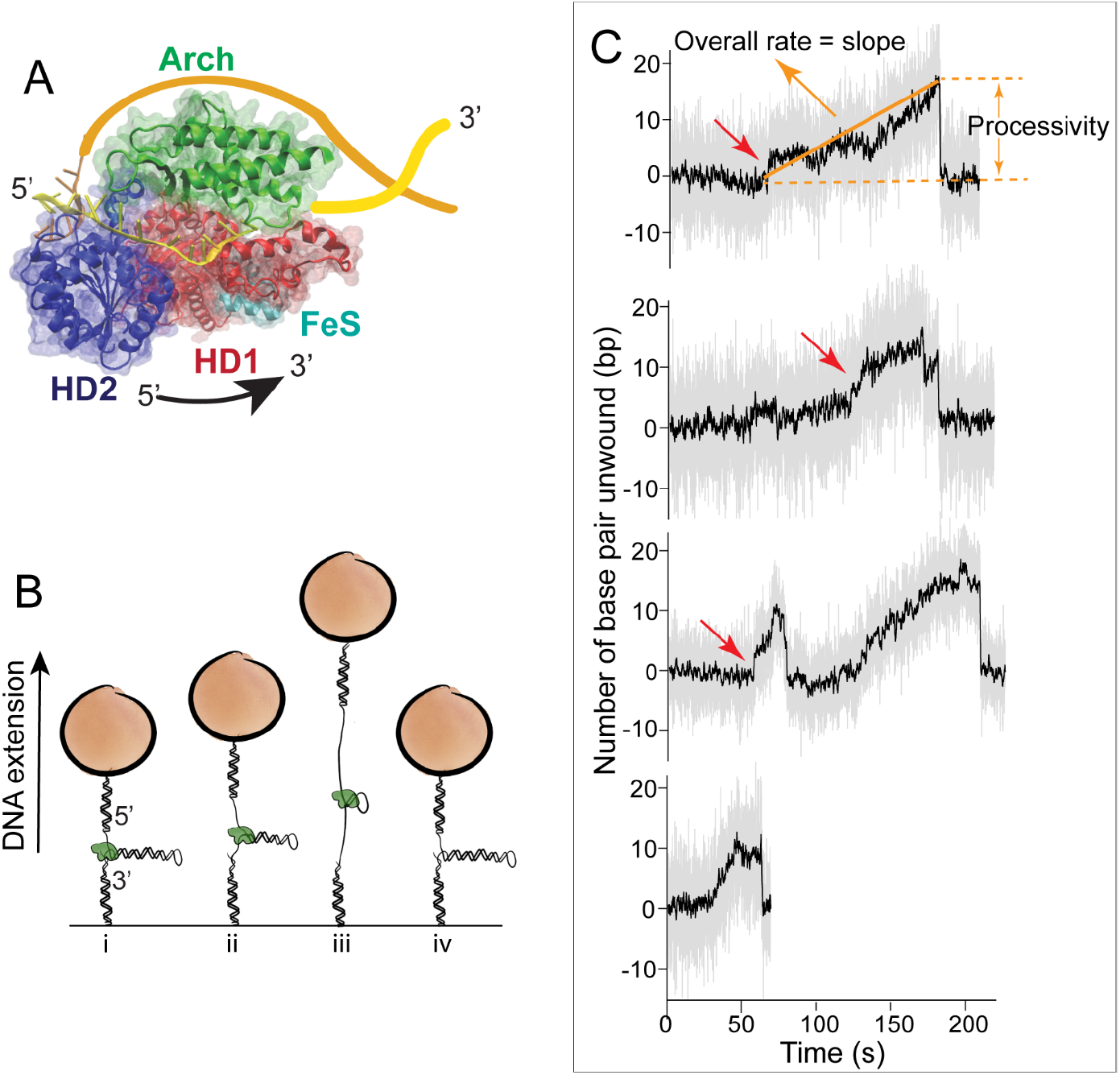
Helicase activity of human XPD (hXPD). A) The structure of XPD with DNA (PDB 6RO4). HD1, HD2, FeS, and Arch domains are labeled. The presumed path of the DNA with respect to XPD is drawn in yellow (translocation strand) and orange (displaced strand). B) Magnetic tweezers assay for helicase activity (not to scale). Helicase activity is detected as an increase in DNA extension as the helicase separates the strands of a 537 bp hairpin (steps i to iii). Dissociation of hXPD from DNA leads to an immediate annealing of two ssDNAs, resulting in a sudden drop in extension (step iv). C) Example helicase extension traces of hXPD. Gray traces are raw data and black lines are Boxcar smoothed traces (window of 31). Fast unwinding runs are indicated by red arrows. Processivity is defined as the maximum number of base pairs unwound in an event (see label on the first trace), and overall unwinding rate as the processivity divided by the total time to reach the maximum (represented by the slope of the orange line on the first trace).

The helicase activity of human XPD within the core TFIIH complex has been measured in both ensemble and single molecule fluorescence experiments (4, 9). However, the basal helicase activity of hXPD is very low in the absence of other TFIIH components such as p34, p44, and p62 (8, 10–13) and has not been studied. Details such as processivity, pausing behavior, and unwinding rate have not been determined. Optical trap measurements have shown that archaeal XPD is a partially active helicase that unwinds dsDNA with single base pair steps and frequent backsteps (14, 15). The ssDNA translocation and conformational dynamics of archaeal XPD have also been studied with single molecule fluorescence methods (16, 17). While these experiments have provided many insights into the archaeal enzyme’s activity, there are likely to be important differences in the activity of the archaeal and human enzymes. The eukaryal and archaeal XPDs are structurally similar, however they only share a 25% sequence identity (18). Archaeal XPD is not part of a TFIIH complex and appears to function on its own. Helicase activities of eukaryotic XPD and TFIIH core complex are inhibited by bulky DNA lesions (19, 20) while conflicting results have been reported for archaeal XPDs (21–23). Additionally archaeal organisms have very different living environments from eukaryotes.

Many mutations in hXPD are the causes of cancers and autosomal recessive disorders such as Trichothiodystrophy (TTD), Cockayne syndrome (CS), and Xeroderma pigmentosum (XP) (24–27). Many of these mutated sites are involved in interactions with binding partners, which are not present in Archaea. Details of basal hXPD activities will provide a basis for better understanding the impact of these interactions on hXPD activity as well as the mechanism of the clinical phenotypes, which will in turn aid in drug design targeting hXPD related diseases (27).

Characterization of hXPD outside of TFIIH is challenging due to its low activity. Here we measured the double stranded DNA unwinding and single stranded DNA translocation activities of hXPD with single molecule magnetic tweezers. Magnetic tweezers offer a robust system for detecting infrequent events due to their stability over time, single molecule resolution, and ability to track multiple molecules at the same time. Our results show that hXPD unwinds DNA with low processivity and slow rate. Comparison of the unwinding and single stranded DNA translocation rates of hXPD reveals that hXPD outside of TFIIH is an intrinsically slow helicase.

## MATERIALS AND METHODS

### Protein Purification

The plasmid carrying human XPD (hXPD) with C-terminal FLAG tag (pFastBac-dual FLAG-XPD_p8) was kindly given to us by Dr. Wei Yang from NIDDK, NIH. The C-terminal FLAG tag has been previously shown to not affect activity in the TFIIH core complex (20). Sf9 cell pellets with XPD expression were purchased from GenScript (NJ, USA). To purify XPD, two grams of cell pellet were suspended in 16 mL of lysis buffer (20 mM TrisCl, pH8, 0.1 mM EDTA, 150 mM KCl, 20% glycerol, 0.1% Triton X-100, 10 mM DTT, 0.25 mM PMSF, 1x protease inhibitor cocktail (EDTA free, 100 x in DMSO, MCE)), pressed at 2500 psi once, and then centrifuged at 200,000 x g for 60 min at 4°C. The supernatant was loaded onto an equilibrated DIY mini-column with DYKDDDDK Fab-Trap™ Agarose resin (250 uL, ffa-10, Chromotek). The column was washed with 6 × 1mL of wash buffer (20 mM TrisCl, pH8, 150 mM KCl, 20% glycerol, 0.01% Triton X-100, 10 mM DTT, 0.25 mM PMSF) and eluted with 4 × 1 mL 0.15 mg/mL (50 µM) 3 x FLAG peptide (Chromotek). The elutant was concentrated with Amico-4 (50k MWCO) and washed with 4 × 1mL concentrating buffer (40 mM TrisCl pH 7.5, 50 mM NaCl,10 mM DTT) at 7500 x g at 4°C. The final XPD protein was aliquoted (20 µL/tube), flash frozen with liquid N2, and stored in concentrating buffer plus 20% glycerol in -80° freezer. Each 20 µL aliquot was further aliquoted into 1 or 2 µL/tube for use and flash frozen with liquid N2. To minimize loss of activity each small aliquot of XPD was used only once to avoid multiple freeze-thaw cycles. Loss of activity was observed without flash freezing. All buffers used during XPD purification were purged under Argon for 30 min to decrease oxygen content. LC-MSMS was used to verify that hXPD was the most abundant protein present with the main contaminants to be heat shock proteins (HSPs). No other TFIIH components were detected.

No direct interactions between HSPs and DNA or TFIIH/XPD have been reported in the literatures and the involvement of HSPs in DNA repair is mainly their chaperone functions (28). Therefore, we do not expect any effects of the contaminant HSPs on our studies. ICP-MS analysis of our hXPD preparation determined the content of Fe in XPD to be ~ 50% of the expected full occupancy, indicating that a population of the enzyme retained the FeS cluster during purification. The total concentration of the protein was estimated from SDS stain gel, and the purity was estimated to be ~70%. To account for the loss of the FeS cluster, we assumed the concentration of active protein is 50% of the estimated total concentration.

### DNA preparation

For magnetic tweezers experiments, the DNA substrate used is a 537-bp hairpin with a short dsDNA handle on 5’ end and long dsDNA handle on 3’ end, the 5’ handle contains a 25-nt single stranded loading region (GTTTGCATTTTGGTTGGTTTCACCG) adjacent to the hairpin (Figure S4). We constructed the hairpin according to the procedures previously described (29) with minor modifications. Complementary strands of DNA handles were first annealed separately by incubating at 94°C for 5 minutes and then gradually cooled to 10° (−1°C/30 s) in 50 mM NaCl and 10 mM TrisCl, pH 7.5. The top handle was modified with biotin on the 5′ end for attachment to a streptavidin coated magnetic bead. The bottom handle was modified with a 5’ overhang (ACCA) on the complementary strand for ligating to the following dsDNA long handle with complementary overhang. The 500-bp dsDNA handle was the PCR product of a segment of pBluescript II KS using primers containing BfuAI and BSaI recognition sequences (forward primer: GCTGGGTCTCGTGGTTTCCCTTTAGTGAGGGTTAATTG; reverse primer: TATAGTCCTGTCGGGTTTCG) in the presence of Digoxigenin-11-dUTP (Roche; dTTP/dUTP = 4.5). The Digoxigenin-modified 500-bp handle DNA was digested to create the complementary overhang, purified with Roche PCR purification kit, and ligated to the hairpin with short handles with T4 DNA ligase (New English Biolabs) to finish the 537-bp hairpin preparation.

### Magnetic tweezers experiments

Flow cells were prepared as described (29) with minor modifications. Briefly, 5–10 × 10^−15^ mol DNA substrate was mixed with ~200 ng anti-digoxigenin antibody (Sigma-Aldrich) in PBS and incubated in a sample cell for one hour at room temperature or overnight at 4° C. Unbound DNA was washed out with 2 × 200 µL PBS and then 100 µL of 2% (w/v) BSA in PBS was added to the flow cell to passivate the surface for 2 hours at room temperature. The flow cell is washed with 2 × 200 µL wash buffer (PBS supplemented with 0.02% v/v Tween-20 and 0.3% w/v BSA) before introducing 1-μm MyOne streptavidin-coated magnetic beads (Invitrogen). After 10 minutes incubation with magnetic beads, the sample cell was washed with 3 × 200 μL wash buffer, followed by 200 μL reaction buffer (25 mM TrisCl, pH 8.0, 50 mM KCl, 2 mM MgCl_2_, 0.3% (w/v) BSA, 5% glycerol, 2 mM DTT).

Measurements were performed on a home-built magnetic tweezers instrument. An average force calibration versus magnet position for magnetic beads was obtained using Brownian motion analysis of 6 kb DNA tethers. In a typical experiment, a force of ~12 pN was applied to the bead with the magnetic tweezers to keep the DNA extended and away from surface without distorting or melting the hairpin. Fields of view with multiple usable tethers were first identified. Tethers were considered usable if the hairpin could be mechanically melted by applying a force of 20 pN. The desired concentration of XPD in reaction buffer with 2 mM ATP was added to the flow cell at ~5 pN. Once equilibrated, the force was set back to 12 pN. The z position of the bead was tracked over time to monitor the changes in extension of DNA tether with occasional switching to 20 pN to open the hairpin for approximately 50 seconds before returning to 12 pN. All data were collected at 60 Hz at room temperature (21 ± 2 °C). The image tracking was accomplished with a modified Labview program (Labview 2019, NI) based on the program of Prof. Michael Poirier of Ohio State University.

### Data analysis

The changes in DNA extension were recorded in nm. For simplicity, they were converted to base pair (bp) before any analyses were done. For this conversion, a 20 pN force would be used to manually open hairpin and the extension change of the open hairpin would be measured. Using this open extension and the known 537 bp in the hairpin, we calculated the conversion factor (bp/nm). This was done for almost all DNA tethers, for a few early traces that conversion factors were not measured, an average conversion factor was used.

To obtain processivity and overall rate, the extension traces were first Boxcar smoothed with a window of 31 and then an Igor procedure was used to measure both (Igor Pro 9, WaveMetrics). To analyze features within events, a student t-test based Python procedure was used to fit Boxcar smoothed (window 200), isolated individual events. Exponential decay was used to fit all distributions except distributions of rates, which we found were best fitted with gamma distribution (Python). The errors reported in the manuscript are the errors of fits unless otherwise indicated.

## RESULTS

### hXPD is a low-processivity and overall slow helicase

We used a well-established magnetic tweezers assay to measure the 5’ to 3’ helicase activity of hXPD (Figure 1B) (30, 31). In these experiments, a 537-bp DNA hairpin was attached to the bottom of a flow cell on 3’ end and a paramagnetic bead on 5’ end *via* dsDNA handles. The 5’ handle contains a 25-base single stranded hXPD loading region adjacent to the hairpin. A force of ~12 pN was applied to the bead with the magnetic tweezers to keep the DNA extended and away from the surface without distorting the hairpin (Figure 1B, step i). The z position of the bead was tracked over time in the presence of hXPD and 2 mM ATP to monitor the changes in extension of DNA tether. Helicase activity was detected as an increase in the extension of DNA as XPD helicase unwound the hairpin, releasing two nucleotides from each base pair unwound (Figure 1B, steps ii and iii). Since the magnetic force applied was not enough to open the DNA hairpin, dissociation of helicase from DNA led to an immediate reannealing of the unwound strands (Figure 1B, step iv). The helicase could bind and start unwinding again; therefore, multiple events could be observed on a single tether over the course of an experiment. Unwinding events that start at the 5’ ssDNA loading region at the base of the hairpin are referred to as “base unwinding” from here on.

Shown in Figure 1C are example extension traces of hXPD base unwinding activity. Typical unwinding events include periods of fast and slow unwinding, pauses, and backsteps. We quantified unwinding processivity from extension traces. Unwinding processivity is defined as maximum number of base pairs unwound before protein dissociates from DNA (distance between two dashed orange lines in the first example trace of Figure 1C). Consistent with the low activity observed in ensemble measurements, the majority of events exhibited low processivity. In many of the events, hXPD stalled at the first four positions of the hairpin before resuming further unwinding (Figure S1). This early unwinding stalling may be the result of high GC content in that region (14). An exponential fit to the processivity data gave a mean of 13 ± 1 bp, similar to previously observed for archaeal XPD (14). No helicase activity was observed in the absence of ATP (compare Figures S2A and S2B).

Overall rates for base unwinding were also obtained from extension traces. For each event the overall unwinding rate was calculated from total number of bases unwound (processivity) divided by the time to reach the maximum (slope of the orange line, Figure 1C). We obtained a mean overall rate of 0.25 ± 0.03 bp/s, indicating that hXPD is a slow helicase relative to other SF2 helicases (14, 32).

The unwinding data were collected at multiple hXPD apparent concentrations ranging from 0.1 nM to 5 nM. We obtained the same mean processivity and overall rate for each concentration (Table S1), indicating we were measuring activities of individual hXPD enzymes. We did not observe higher processivity with increasing hXPD concentration as reported for archaeal XPD, which the authors attributed to multiple XPD binding (14). This may be because the actual concentrations of active hXPD in our experiments were lower than the apparent concentrations and the probability of binding two or more hXPD to the 25-nt loading region was low at the concentration range used here. We cannot be sure of the exact concentration of active helicase in our experiments due to instability of the iron sulfur cluster. The iron sulfur cluster is known to be essential for activity and is easily lost due to oxidation (33–35). The fact that we observed activity at all indicated that at least some population of the hXPD in our experiments retained the FeS cluster, as verified by the ICP-MS analysis. We also measured time intervals between two measurable, consecutive events at several XPD concentrations and found that the intervals decrease with increasing hXPD concentration (Table S1, Figure S3), consistent with more hXPD binding at higher concentrations.

### Unwinding is a reflection of DNA translocation

#### Middle unwinding and translocation with refolding

To further characterize the activity of hXPD, we manually opened the hairpin DNA under 20 pN force in the presence of hXPD and 2mM ATP and held it open for ~50 seconds. Doing so created a ~1 kb ssDNA on which the enzyme could bind at any position (Figures 2A and 2C, step ii). Binding of hXPD to the newly exposed ssDNA region could block hairpin refolding when the force was returned to 12 pN (Figures 2A and 2C, step iii). From there two scenarios were possible. If hXPD was bound at the 5’ end of the ssDNA-dsDNA junction of the partially folded hairpin it could initiate unwinding (which we defined as “middle unwinding”, Figure 2A, steps iii and iv). If hXPD was bound at 3’ end of the ssDNA-dsDNA junction it could translocate on the remaining ssDNA in the 5’-3’ direction. In this scenario the hairpin would close behind the enzyme as it translocated on the single stranded DNA, giving a measurement of XPD translocation as the hairpin refolded behind it (referred to from here on as translocation with refolding, Figure 2C, steps iii and iv).

**Figure 2.**
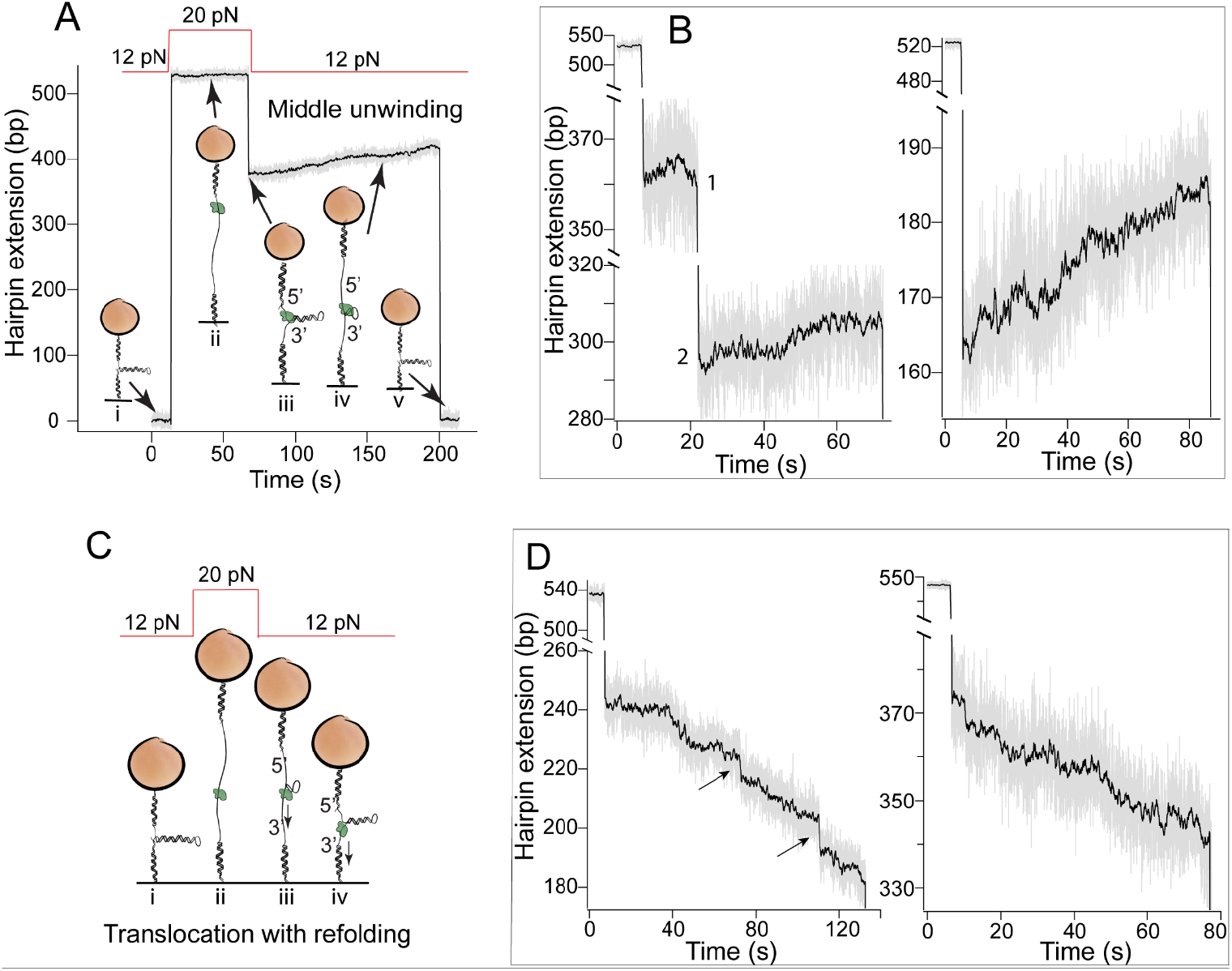
Middle unwinding and translocation with refolding of hXPD. A) Experimental design for middle unwinding (not to scale). Hairpin was opened at high force (20 pN) in the presence of hXPD, allowing hXPD binding to the newly exposed ssDNA region (step ii). When force was returned to 12 pN to permit re-annealing of the hairpin strands, a 5’ end-bound hXPD could block the re-annealing (step iii) and start unwinding the partially annealed hairpin (step iv). Dissociation of hXPD leads to sudden drop in extension (step v). B) Example extension traces of hXPD middle unwinding. Separate unwinding events in the first trace are labeled as 1 and 2. C) Experimental design for translocation with refolding (not to scale). After the hairpin was opened (20 pN, step ii) and force returned to 12 pN, a 3’ end-bound hXPD could block the re-annealing (step iii) and start translocating along the ssDNA with hairpin re-annealing behind it (step iv). Arrow shows direction of helicase translocation. D) Example extension traces of hXPD translocation with refolding. Gray traces are raw data. Black lines are Boxcar smoothed traces (window of 31). Arrows on the first trace indicate “sliding” activity which we interpret as distinct from translocation with refolding.

Examples of hXPD middle unwinding after melting and partial refolding of the hairpin are shown in Figures 2A and 2B. We determined the processivity (16 ± 1 bp) and overall rate (0.34 ± 0.06 bp/s) in the same manner as for the base unwinding. In this case, hXPD could explore the whole 537-nt 5’ half of the unfolded hairpin, which provided a randomized sequence (sequence shown in Figure S4). The similar processivity to that of the base unwinding indicates that the low processivity at the base of the hairpin is not due to the sequence of that region. Since both middle and base unwinding represented the same activity, we combined the data into a single set (duplex unwinding, Figures 3A and 3B). The mean processivity and overall rate of the duplex unwinding are 14 ± 1 bp and 0.28 ± 0.03 bp/s, respectively (Table 1).

**Table 1.** Mean processivity and overall rate

| **Activity** | **Processivity<sup>a</sup>**<br>**(bases)** | **Overall**<br>**rate<sup>b</sup>**<br>**(base/s)** | **No. of events**<br>**(processivity/**<br>**rate)** | **No. of**<br>**Tethers** | **No. of**<br>**flow cells** |
| --- | --- | --- | --- | --- | --- |
| Duplex<sup>c</sup><br>unwinding | 14 ± 1 | 0.28 ± 0.03 | 900/829 | 115 | 27 |
| Translocation<br>with refolding | 44 ± 1 | 0.40 ± 0.06 | 1025/1007 | 150 | 22 |
| ssDNA<br>translocation | NA | 0.26 ± 0.09 | NA/358 | 110 | 21 |

Means and errors are from exponential decay<sup>a</sup> or gamma distribution<sup>b</sup> fits.
<sup>c</sup>Duplex unwinding is the combination of base and middle unwinding.

**Figure 3.**
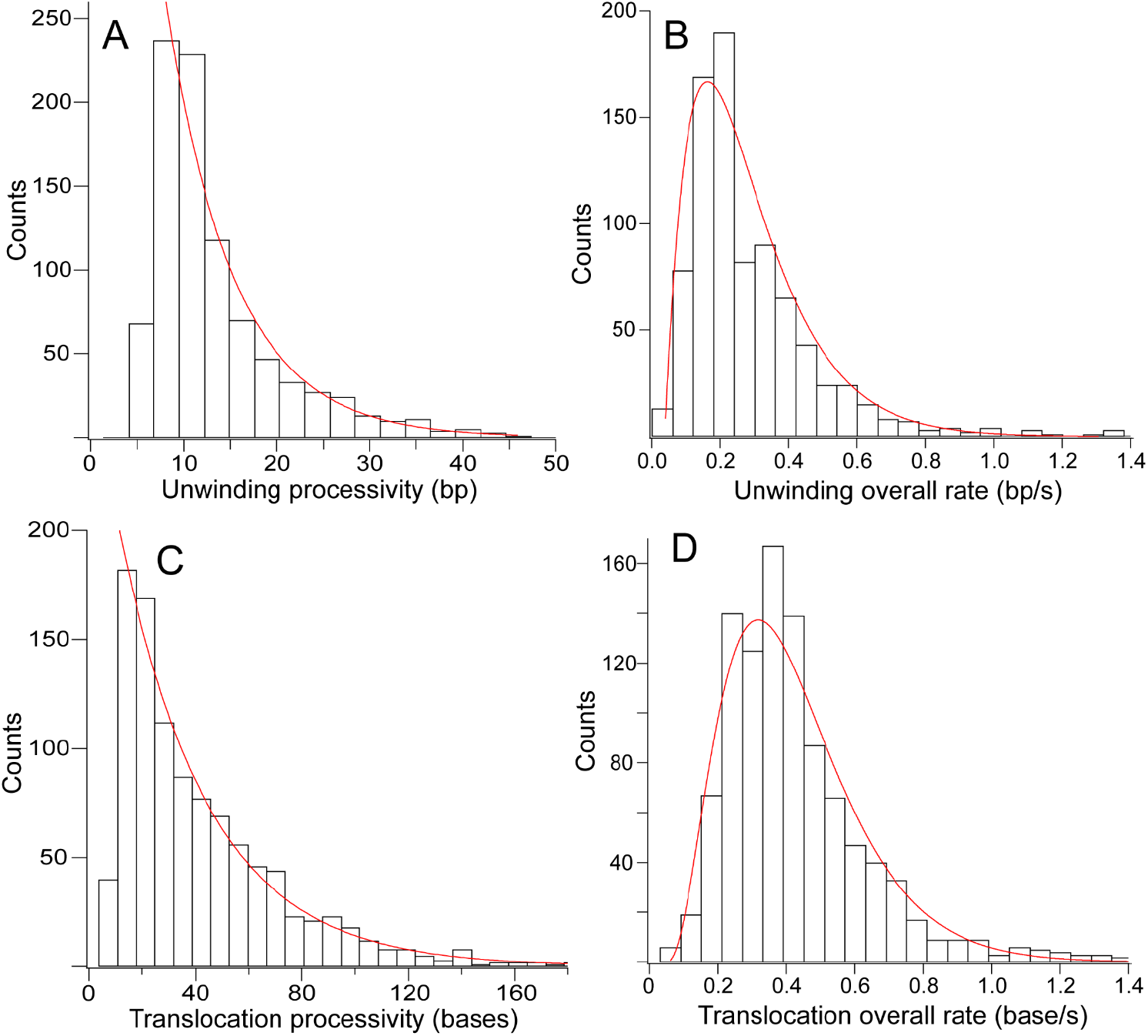
Histograms of processivities and overall rates for hXPD duplex unwinding and translocation with refolding. A) Processivity of hXPD duplex unwinding. Duplex unwinding includes both base and middle unwinding. B) Overall duplex unwinding rate. C) Processivity of hXPD translocation with refolding. D) hXPD overall rate for translocation with refolding. Red curves are exponential decay or gamma distribution fits of the data.

Figure 2D shows the example traces of hXPD translocation with refolding. Analysis of translocation traces gave a mean processivity of 44 ± 1 bases and a mean overall rate of 0.40 ± 0.06 base/s (Figures 3C and 3D, Table 1). This rate indicates that the overall slow unwinding of hXPD is mainly due to slow translocation, rather than the result of contributions from base pairing in the duplex. The much higher mean processivity (compared to duplex unwinding) and broader distribution (Figure 3C) suggest either a higher processivity for translocation with refolding or/and more than one events unintentionally counted as one during our analysis. Due to the limitations of our experiments, we cannot distinguish between the two possibilities. We regarded abrupt drops of 5 or more bases as the end of an event in our processivity analysis (Figure 2D arrows). Events 4 bases or less apart from each other would have been counted as one event. These events could be the results of partial dissociation then re-engaging of a protein with the DNA. In the case of unwinding the duplex re-annealing force might prevent reengagement, limiting the processivity. Binding and sliding, but not translocation, of XPD on ssDNA was observed in the absence of ATP (compare Figures S2C and S2D).

#### Analysis of hXPD unwinding and translocation with refolding events

The overall duplex unwinding rate of ~ 0.28 bp/s (Table 1) is slow compared to previously measured rates of ~10 - 20 bp/s for archaeal XPD (14, 17) and the human core TFIIH complex (9). However, we frequently observed different behaviors within individual events, including fast runs and pauses (Figure 1C). To better understand these behaviors, we analyzed the features within individual events. Extension traces were Boxcar smoothed with a window of 200, then individual events were isolated (ending at the maximum of the event) and fitted with a student t-test based Python procedure to obtain pause times, run sizes, and run rates (Figure 4A). The window size of 200 for smoothing was chosen to preserve the main features of an event. For the purpose of this analysis, any runs less than 2 bp were not considered.

**Figure 4.**
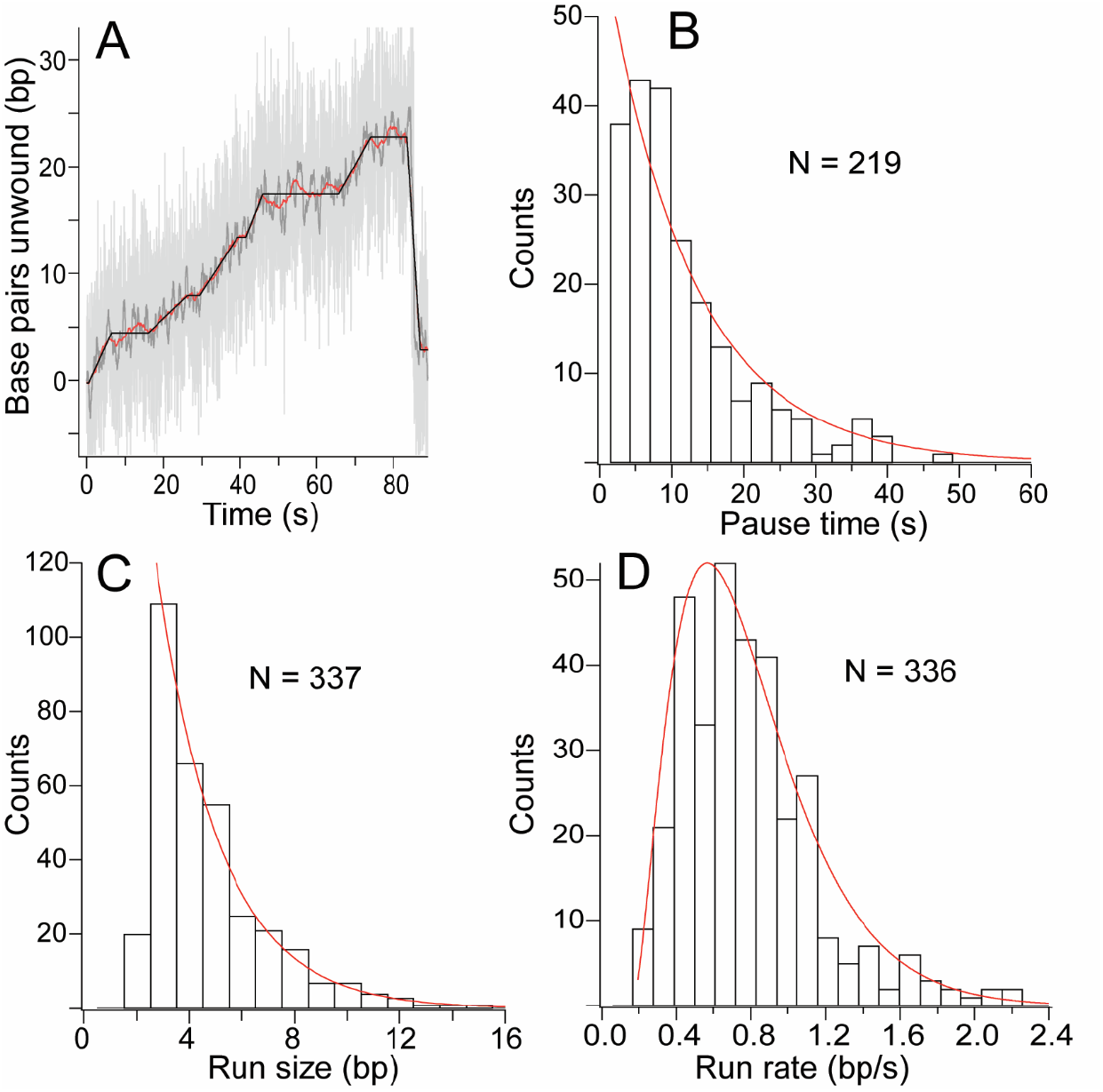
Analysis of individual unwinding events. A) Example of unwinding event with different windows of Boxcar smoothing and t-test based step finding fit. Light grey: raw data; Dark grey: smoothed data with a window of 31; Red: smoothed data with a window of 200; Black: t-test based fit. B) Histogram of pause times fitted with exponential curve. C) Histogram of run sizes fitted with exponential curve. D) Histogram of run rates with gamma distribution fit.

The pauses obtained from the analysis represented stalled states of the enzyme, as some short pauses were lost during smoothing. Therefore, some pauses may include repetitive short forward and backward steps which did not lead to measurable unwinding. The distribution of pause times between runs is shown in Figure 4B. The mean pause from the exponential fit was 13 ± 1 s (Table 2). Figure 4C shows the run size distribution. An exponential fit to the data gave a mean size of 4.9 ± 0.1 bp (Table 2). We observed a small number of backsteps greater than our 2-bp cutoff. The fundamental steps are expected to be one base (14); therefore, our fitted runs consisted of multiple fundamental steps. Analysis of the run rate distribution gave a mean run rate of 0.7 ± 0.2 bp/s, with a small population of rates more than 1.2 bp/s (Figure 4D). The high rates here were underestimated due to smoothing.

**Table 2.** Parameters from analysis of individual events

| **Activity** | **Pause time<sup>d</sup>**<br>**(s)** | **Run size<sup>d</sup>**<br>**(base)** | **Run rate<sup>e</sup>**<br>**(base/s)** |
| --- | --- | --- | --- |
| Duplex<br>unwinding | 13 ± 1 | 4.9 ± 0.1 | 0.7 ± 0.2 |
| Translocation<br>with refolding | 9 ± 1 | 6.1 ± 0.4 | 0.8 ± 0.3 |

Means and errors are from exponential decay<sup>d</sup> or gamma distribution<sup>e</sup> fits.

To more accurately determine the rate of fast runs, we analyzed unwinding traces with less smoothing (window of 31). We found a subset of runs with rates 3-10 bp/s (N = 80). Although the mean run rate is faster than the overall rate of hXPD, it is still very slow compared to the overall rate of ~10 - 20 bp/s previously measured (9, 14, 17). We also compared the total time the enzyme spent stalling to the total time spent unwinding from all events and found them to be approximately the same, revealing the contribution of pauses to the low unwinding rate in addition to the already slow translocation.

We also analyzed the events from hXPD translocation with refolding (Figure S5). We obtained the mean pause time of 9 ± 1 s. The shorter pause time is consistent with the slightly higher overall rate and may be the result of the DNA annealing force behind hXPD. The run size (6.1 ± 0.4 bp) and run rate (0.8 ± 0.3 bp/s) were similar to those of duplex unwinding (Table 2). The similarity in run size and rate to those of unwinding confirms that unwinding is a reflection of the hXPD translocation activity.

### Sequence affects initiation positions of hXPD unwinding but not translocation

To investigate potential DNA sequence dependence of hXPD activity, the initiation positions of middle unwinding and translocation with refolding on the unfolded hairpin were obtained and histogrammed (Figure 5). The distribution of the unwinding initiation positions (Figure 5A) showed that hXPD could bind and start unwinding at most regions on the 537-nt 5’ region of the unfolded hairpin. However, there were two regions with minimal initiation for unwinding (from ~ 110 to 150 bp and ~350 to 430 bp). The only distinct feature of the regions is the presence of the poly-A tracts (see Figure S4 for sequence, highlighted in blue). Poly-A tracts are known to give rise to unique structures and properties (36, 37), which could affect hXPD activity. We also observed one location where initiation was slightly enhanced (~ 200 bp). This region contains 3 GC stretches clustered together, followed by AT-rich regions. However, we do not have the resolution to determine the exact location on the hairpin. The plots of processivity (Figure S6A) or overall rate (Figure S6B) *vs*. initiation positions of middle unwinding did not show a strong sequence dependence. However, there were a small number of higher processivity and rate outliers at the same region where enhanced initiation occurred. For translocation with refolding, no sequence dependence of translocation initiation was observed (Figure 5B). We note that the sequence that hXPD translocated on was the complementary strand of the sequence shown in Figure S4.

**Figure 5.**
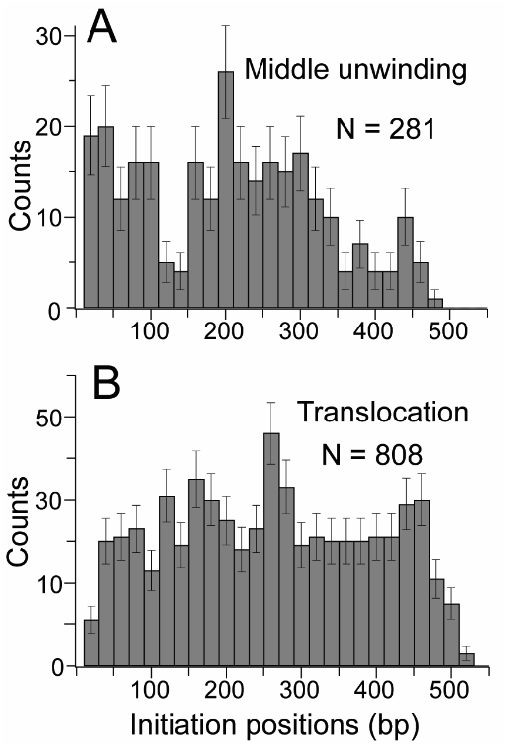
Histograms of hXPD initiation positions on unfolded hairpin for middle unwinding (A) and translocation with refolding (B).

### Displaced strand has no detectable effect on rate

To test the effect of the displaced strand on unwinding rate, we opened the hairpin under high force (20 pN) during an active unwinding event (Figure 6A, steps ii and iii) and held it open for ~50 seconds. During this time hXPD could switch from unwinding to translocating on newly exposed ssDNA (Figure 6A, step iii). After 50 s we reduced the force to 12 pN, allowing the two ssDNA to partially re-anneal and hXPD to switch back to unwinding (Figure 6A, steps iv and v). When opening the hairpin with force during duplex unwinding, we eliminate the interaction between the displaced strand and the Arch domain (8). We also eliminate any opposing re-annealing force. The translocation events in Figure 6A step iii therefore represent true ssDNA translocation. During hXPD translocation with refolding, the two ssDNAs anneal behind hXPD as it walks along ssDNA towards 3’ end. The annealing force could push the protein from behind, increasing the observed translocation rate. To eliminate the effects of a potential assisting re-annealing force, we also opened the hairpin during events of translocation with refolding to measure unassisted ssDNA translocation. The two ssDNA translocation cases created during unwinding and translocation with refolding represented the same activity and were combined into one data set (ssDNA translocation).

**Figure 6.**
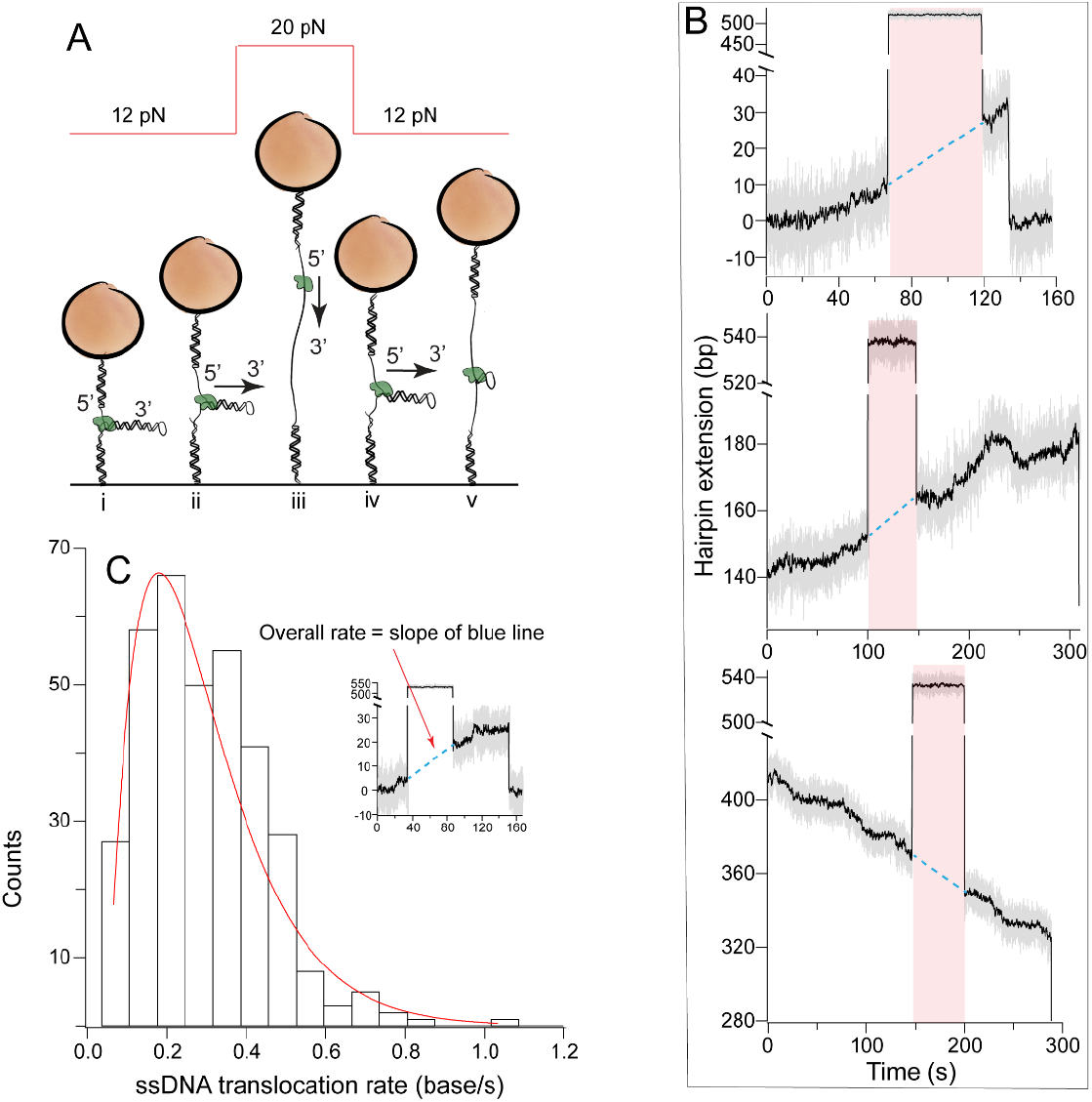
Measurement of ssDNA translocation. A) Experimental design (not to scale) for ssDNA translocation during unwinding. Hairpin was opened at high force (20 pN) for ~50 s (step iii) during an active helicase event (step ii). The helicase continues translocating on the ssDNA (step iii, ssDNA translocation). When the force was returned to 12 pN to permit reannealing of the hairpin strands (step iv), the 5’ end-bound hXPD resumes unwinding the partially annealed hairpin (steps iv and v). B) Example extension traces of ssDNA translocation of hXPD during base unwinding, middle unwinding, and translocation with refolding. Gray traces are raw data and black ones are Boxcar smoothed traces (window of 31). Pink shading indicates duration of ssDNA translocation and blue dashed line represents presumed continuation of helicase translocation on ssDNA. C) Histogram of hXPD overall ssDNA translocation rate with gamma distribution fit (red curve). Inset: overall ssDNA translocation rate is the slope of the dashed blue line.

We see from example traces in Figure 6B that hXPD could continue translocating on ssDNA with a similar rate as prior unwinding or translocation when the hairpin was open. The helicase resumed unwinding or translocating when the force was reduced to allow re-annealing. The distribution of ssDNA translocation rates is shown in Figure 6C. The ssDNA translocation rate is defined as the difference in DNA extension immediately before and after opening divided by the duration of hairpin opening (Figure 6C inset, slope of the blue dashed line). The mean rate for ssDNA translocation is 0.26 ± 0.09 bp/s (Table 1), similar to that of dsDNA unwinding (0.28 ± 0.03 bp/s), indicating no measurable effect on the rate from displaced strand under our experimental setup. This ssDNA translocation rate demonstrates that the slow rate is an intrinsic property of the hXPD. While our observations of the modest sequence dependence above suggest that base pairing energy affects unwinding, the contribution to the rate appears to be minimal.

## DISCUSSION

Our results reveal that human XPD has low processivity (~14 bp) and slow overall rate (~0.3 bp/s, Table 1). However, we observed fast and slow runs and pauses within events. The presence of different rates within events suggests that there might be two modes of hXPD: in one hXPD binds DNA tightly producing fast unwinding runs with fewer backsteps and shorter pauses and in the other hXPD binds less tightly, leading to more backsteps and longer pauses. Incorporation of hXPD into the TFIIH core complex may bias it towards the tighter binding mode, leading to a higher overall rate, as shown in single molecule fluorescence studies of TFIIH where authors observed complete unwinding of a 18-bp dsDNA with no obvious backstepping or pausing (9).

We observed frequent stalling at the first four bases of the hairpin, corresponding to a region with high GC content. This suggests that the enzyme’s activity is affected by local duplex properties. In agreement with this we also observed a modest sequence dependence for initiation positions of unwinding on the unfolded hairpin, but not for those of translocation with refolding. However, we have shown that under our experimental conditions the displaced strand has no detectable effect on the rate of hXPD unwinding. It has been reported that a significant percent of base pairs at the ds/ssDNA fork are open at any given time at room temperature due to thermal breathing (38). The slow translocation of hXPD and frequent thermal breathing of base pairs at the DNA fork could diminish the effect of DNA sequence on unwinding. In addition, our DNA hairpin was not optimized to maximize the effect of sequence, which further limits our ability to observe the effect of sequence dependence. With these factors in mind, we propose that hXPD is most likely a partially active helicase, like its archaeal counterpart (14).

Our results represent the first direct measurement of the DNA unwinding and ssDNA translocation of human XPD by itself. Human XPD is dynamic and has multiple roles in the cell. The nucleus must maintain a balance of both free XPD and TFIIH bound XPD (39). Therefore, the cell needs to regulate the helicase activity to accommodate the fluctuations in damage response. The observations of the much lower ssDNA translocation and unwinding rates than those for the archaeal XPD and human core TFIIH complex suggest that other binding partners in the cell regulate the rate of hXPD helicase activity. The measurements presented here provide a baseline for future studies of the effects of TFIIH components such as p34, p44, p62, and XPB on hXPD activity.

## AUTHOR CONTRIBUTIONS

C. Mao: designed and performed experiments, analyzed data, and wrote the manuscript. M. Mills: designed experiments and wrote manuscript.

## ACKNOWLEDGEMENTS

We would like to thank Professor Gavin King of the University of Missouri for his suggestions and discussions throughout the project, Professor Michael Poirier and Ariel Robbins of Ohio State University for kindly sharing Labview program and help during program adaptation, Dillon Balthrop for sharing the Python programs, Deepesh Sigdel, Dillon Balthrop, and Connor Nance for help with various things in the lab, and Samuel Cheslik and Doniven Hicks for help with flow cell assembly. This work was supported by the National Institutes of Health (5K22HL142846 to M. M.) and the University of Missouri (startup fund to M. M.). Funding for open access charge: National Institutes of Health.

## REFERENCES

1. Ito, S., L.J. Tan, D. Andoh, T. Narita, M. Seki, Y. Hirano, K. Narita, I. Kuraoka, Y. Hiraoka, and K. Tanaka. 2010. MMXD, a TFIIH-independent XPD-MMS19 protein complex involved in chromosome segregation. Mol Cell. 39:632–640.

2. Houten, B.V., J. Kuper, and C. Kisker. 2016. Role of XPD in cellular functions: To TFIIH and beyond. DNA Repair (Amst). 44:136–142.

3. Compe, E., E. Pangou, N. Le May, C. Elly, C. Braun, J.-H. Hwang, F. Coin, I. Sumara, K.-W. Choi, and J.-M. Egly. 2022. Phosphorylation of XPD drives its mitotic role independently of its DNA repair and transcription functions. Sci Adv. 8:eabp9457.

4. Kokic, G., A. Chernev, D. Tegunov, C. Dienemann, H. Urlaub, and P. Cramer. 2019. Structural basis of TFIIH activation for nucleotide excision repair. Nat Commun. 10:2885.

5. Kuper, J., S.C. Wolski, G. Michels, and C. Kisker. 2012. Functional and structural studies of the nucleotide excision repair helicase XPD suggest a polarity for DNA translocation. EMBO J. 31:494–502.

6. Lainé, J., V. Mocquet, and J. Egly. 2006. TFIIH Enzymatic Activities in Transcription and Nucleotide Excision Repair. In: Methods in Enzymology. Academic Press. pp. 246–263.

7. Coin, F., V. Oksenych, and J.-M. Egly. 2007. Distinct Roles for the XPB/p52 and XPD/p44 Subcomplexes of TFIIH in Damaged DNA Opening during Nucleotide Excision Repair. Mol Cell. 26:245–256.

8. Peissert, S., F. Sauer, D.B. Grabarczyk, C. Braun, G. Sander, A. Poterszman, J.-M. Egly, J. Kuper, and C. Kisker. 2020. In TFIIH the Arch domain of XPD is mechanistically essential for transcription and DNA repair. Nat Commun. 11:1667.

9. Bralić, A., M. Tehseen, M.A. Sobhy, C.-L. Tsai, L. Alhudhali, G. Yi, J. Yu, C. Yan, I. Ivanov, S.E. Tsutakawa, J.A. Tainer, and S.M. Hamdan. 2022. A scanning-to-incision switch in TFIIH-XPG induced by DNA damage licenses nucleotide excision repair. Nucleic Acids Res. gkac1095.

10. Coin, F., J.-C. Marinoni, C. Rodolfo, S. Fribourg, A.M. Pedrini, and J.-M. Egly. 1998. Mutations in the XPD helicase gene result in XP and TTD phenotypes, preventing interaction between XPD and the p44 subunit of TFIIH. Nat Genet. 20:184–188.

11. Kuper, J., C. Braun, A. Elias, G. Michels, F. Sauer, D.R. Schmitt, A. Poterszman, J.-M. Egly, and C. Kisker. 2014. In TFIIH, XPD Helicase Is Exclusively Devoted to DNA Repair. PLoS Biol. 12:e1001954.

12. Barnett, J.T., J. Kuper, W. Koelmel, C. Kisker, and N.M. Kad. 2020. The TFIIH subunits p44/p62 act as a damage sensor during nucleotide excision repair. Nucleic Acids Res. 48:12689–12696.

13. Radu, L., E. Schoenwetter, C. Braun, J. Marcoux, W. Koelmel, D.R. Schmitt, J. Kuper, S. Cianférani, J.M. Egly, A. Poterszman, and C. Kisker. 2017. The intricate network between the p34 and p44 subunits is central to the activity of the transcription/DNA repair factor TFIIH. Nucleic Acids Res. 45:10872–10883.

14. Qi, Z., R.A. Pugh, M. Spies, and Y.R. Chemla. 2013. Sequence-dependent base pair stepping dynamics in XPD helicase unwinding. eLife. 2:e00334.

15. Stekas, B., S. Yeo, A. Troitskaia, M. Honda, S. Sho, M. Spies, and Y.R. Chemla. 2021. Switch-like control of helicase processivity by single-stranded DNA binding protein. eLife.

16. Ghoneim, M., and M. Spies. 2014. Direct Correlation of DNA Binding and Single Protein Domain Motion via Dual Illumination Fluorescence Microscopy. Nano Lett. 14:5920–5931.

17. Honda, M., J. Park, R.A. Pugh, T. Ha, and M. Spies. 2009. Single-Molecule Analysis Reveals Differential Effect of ssDNA-Binding Proteins on DNA Translocation by XPD Helicase. Molecular Cell. 35:694–703.

18. Liu, H., J. Rudolf, K.A. Johnson, S.A. McMahon, M. Oke, L. Carter, A.-M. McRobbie, S.E. Brown, J.H. Naismith, and M.F. White. 2008. Structure of the DNA repair helicase XPD. Cell. 133:801–812.

19. Naegeli, H., P. Modrich, and E.C. Friedberg. 1993. The DNA helicase activities of Rad3 protein of Saccharomyces cerevisiae and helicase II of Escherichia coli are differentially inhibited by covalent and noncovalent DNA modifications. J Biol Chem. 268:10386–10392.

20. Li, C.-L., F.M. Golebiowski, Y. Onishi, N.L. Samara, K. Sugasawa, and W. Yang. 2015. Tripartite DNA Lesion Recognition and Verification by XPC, TFIIH, and XPA in Nucleotide Excision Repair. Mol Cell. 59:1025–1034.

21. Mathieu, N., N. Kaczmarek, P. Rüthemann, A. Luch, and H. Naegeli. 2013. DNA Quality Control by a Lesion Sensor Pocket of the Xeroderma Pigmentosum Group D Helicase Subunit of TFIIH. Curr Biol. 23:204–212.

22. Rudolf, J., C. Rouillon, U. Schwarz-Linek, and M.F. White. 2010. The helicase XPD unwinds bubble structures and is not stalled by DNA lesions removed by the nucleotide excision repair pathway. Nucleic Acids Res. 38:931–941.

23. Mathieu, N., N. Kaczmarek, and H. Naegeli. 2010. Strand- and site-specific DNA lesion demarcation by the xeroderma pigmentosum group D helicase. Proc Natl Acad Sci U S A. 107:17545–17550.

24. Abdulrahman, W., I. Iltis, L. Radu, C. Braun, A. Maglott-Roth, C. Giraudon, J.-M. Egly, and A. Poterszman. 2013. ARCH domain of XPD, an anchoring platform for CAK that conditions TFIIH DNA repair and transcription activities. Proc Natl Acad Sci USA. 110:E633–E642.

25. Lehmann, A.R. 2001. The xeroderma pigmentosum group D (XPD) gene: one gene, two functions, three diseases. Genes Dev. 15:15–23.

26. Greber, B.J., D.B. Toso, J. Fang, and E. Nogales. 2019. The complete structure of the human TFIIH core complex. eLife.

27. Fuss, J.O., and J.A. Tainer. 2011. XPB and XPD helicases in TFIIH orchestrate DNA duplex opening and damage verification to coordinate repair with transcription and cell cycle via CAK kinase. DNA Repair (Amst). 10:697–713.

28. Sottile, M.L., and S.B. Nadin. 2018. Heat shock proteins and DNA repair mechanisms: an updated overview. Cell Stress Chaperones. 23:303–315.

29. Seol, Y., M.-P. Strub, and K.C. Neuman. 2016. Single molecule measurements of DNA helicase activity with magnetic tweezers and t-test based step-finding analysis. Methods. 105:119–127.

30. Mills, M., Y.-C. Tse-Dinh, and K.C. Neuman. 2018. Direct Observation of Topoisomerase IA Gate Dynamics. Nat Struct Mol Biol. 25:1111–1118.

31. Seol, Y., and K.C. Neuman. 2011. Magnetic tweezers for single-molecule manipulation. Methods Mol Biol. 783:265–293.

32. Seol, Y., G.M. Harami, M. Kovács, and K.C. Neuman. 2019. Homology sensing via non-linear amplification of sequence-dependent pausing by RecQ helicase. eLife. 8:e45909.

33. Rudolf, J., V. Makrantoni, W.J. Ingledew, M.J.R. Stark, and M.F. White. 2006. The DNA Repair Helicases XPD and FancJ Have Essential Iron-Sulfur Domains. Molecular Cell. 23:801–808.

34. Fan, L., J.O. Fuss, Q.J. Cheng, A.S. Arvai, M. Hammel, V.A. Roberts, P.K. Cooper, and J.A. Tainer. 2008. XPD helicase structures and activities: insights into the cancer and aging phenotypes from XPD mutations. Cell. 133:789–800.

35. Pugh, R.A., M. Honda, H. Leesley, A. Thomas, Y. Lin, M.J. Nilges, I.K.O. Cann, and M. Spies. 2008. The iron-containing domain is essential in Rad3 helicases for coupling of ATP hydrolysis to DNA translocation and for targeting the helicase to the single-stranded DNA-double-stranded DNA junction. J Biol Chem. 283:1732–1743.

36. Haran, T.E., and U. Mohanty. 2009. The unique structure of A-tracts and intrinsic DNA bending. Quarterly Reviews of Biophysics. 42:41–81.

37. Dršata, T., N. Špačková, P. Jurečka, M. Zgarbová, J. Šponer, and F. Lankaš. 2014. Mechanical properties of symmetric and asymmetric DNA A-tracts: implications for looping and nucleosome positioning. Nucleic Acids Research. 42:7383–7394.

38. Jose, D., K. Datta, N.P. Johnson, and P.H. von Hippel. 2009. Spectroscopic studies of position-specific DNA “breathing” fluctuations at replication forks and primer-template junctions. Proceedings of the National Academy of Sciences. 106:4231–4236.

39. Houten, B.V., J. Kuper, and C. Kisker. 2016. Role of XPD in cellular functions: To TFIIH and beyond. DNA Repair (Amst). 44:136–142.

