## Supplemental for "Characterization of human XPD helicase activity with Single Molecule Magnetic Tweezers"

Table S1. Mean processivity, mean overall rate, and interval between events for base unwinding at different hXPD concentrations

| **XPD Concentration** | 100 to 250 pM | 500 pM | 1 nM | 1.5 nM | 2.5 to 5 nM |
| --- | --- | --- | --- | --- | --- |
| **Processivity (bp)^a^** | 12 ± 1 | 12 ± 1 | 12 ± 1 | 12 ± 1 | 14 ± 1 |
| **Overall rate^b^ (bp/s)** | 0.26 ± 0.10 | 0.22 ± 0.11 | 0.29 ± 0.07 | 0.23 ± 0.07 | 0.23 ± 0.09 |
| **Interval between events^c^ (s)** | 4389 ± 953 | 2649 ± 354 | NA | 1327 ± 169 | 1208 ± 292 |
| **No. of events (processivity/rate)** | 86/64 | 157/123 | 136/108 | 170/141 | 76/64 |
| **No. of tethers** | 21 | 27 | 36 | 14 | 17 |
| **No. of flow cells** | 5 | 7 | 6 | 2 | 5 |

Means and Errors are from exponential^a^ and gamma distribution^b^ fits.

**^c^**Means of intervals between two measurable, consecutive base unwinding events are calculated using Wave Stats function from Igor, and errors are standard error of means.

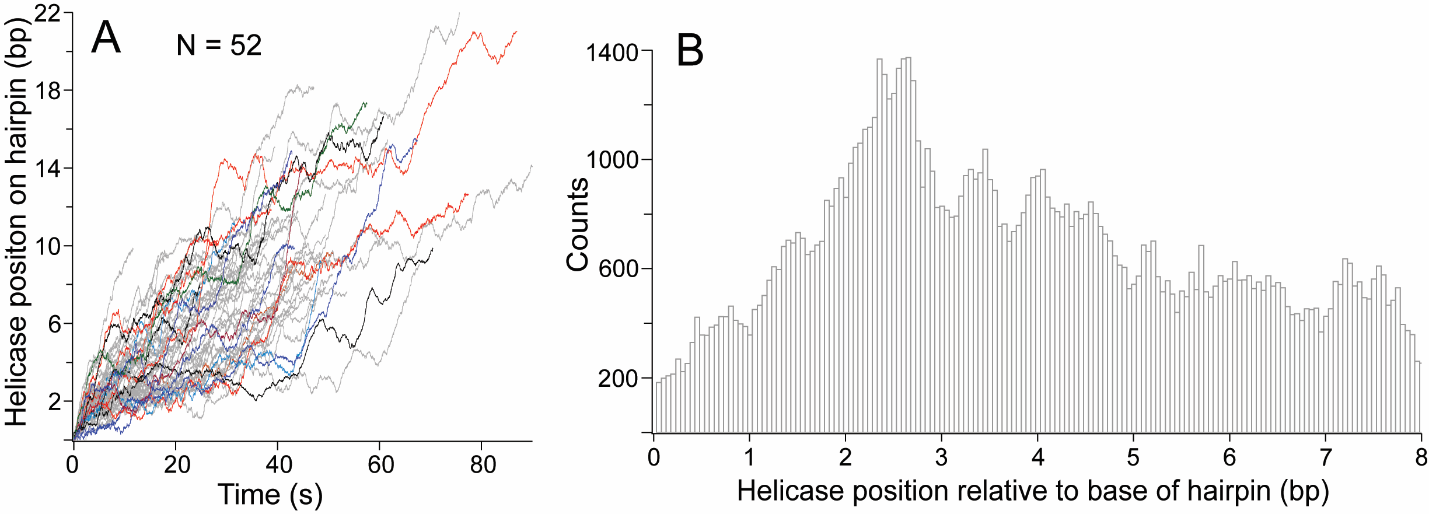

**Figure S1.** Frequent stalling at the positions 2 to 4 of hairpin within base unwinding events. A) Aligned base unwinding events with processivity of 8 bp or more. Base unwinding traces were first Boxcar smoothed with a 200 window, then individual events were isolated and aligned at the beginning of each event (t = 0). Several events are highlighted in color. B) Histogram of helicase positions of aligned events in A.

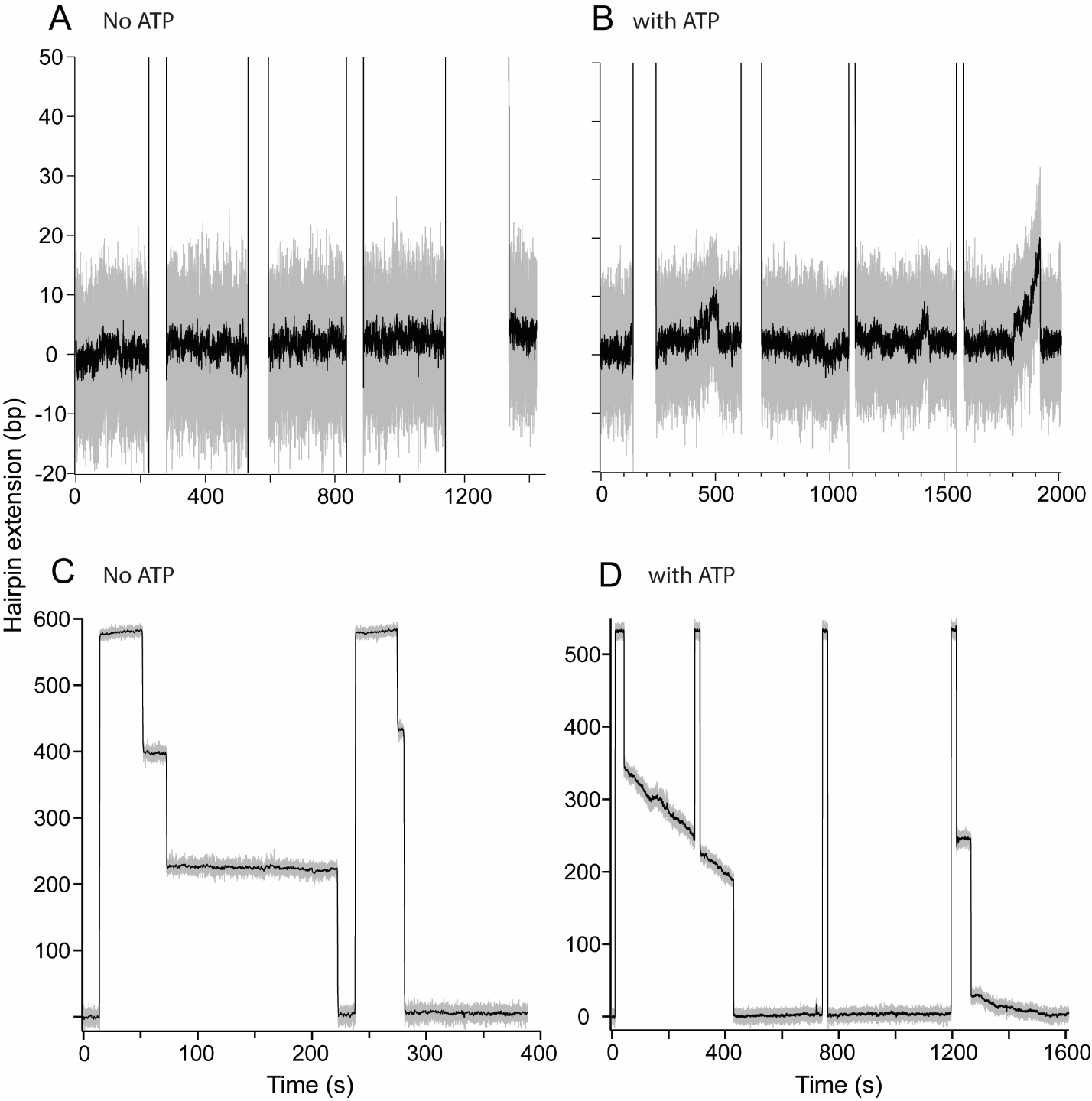

**Figure S2.** Comparison of no ATP control and activity with ATP. A) Typical baseline trace in absence of ATP. B) Unwinding activity in presence of 2mM ATP. C) hXPD binding to ssDNA in absence of ATP. D) ssDNA translocation and binding of hXPD in presence of 2 mM ATP. The gaps in the middle of traces of A) and B) are durations when hairpin was opened under force. Gray traces are raw data. Black lines are Boxcar smoothed traces (window of 31).

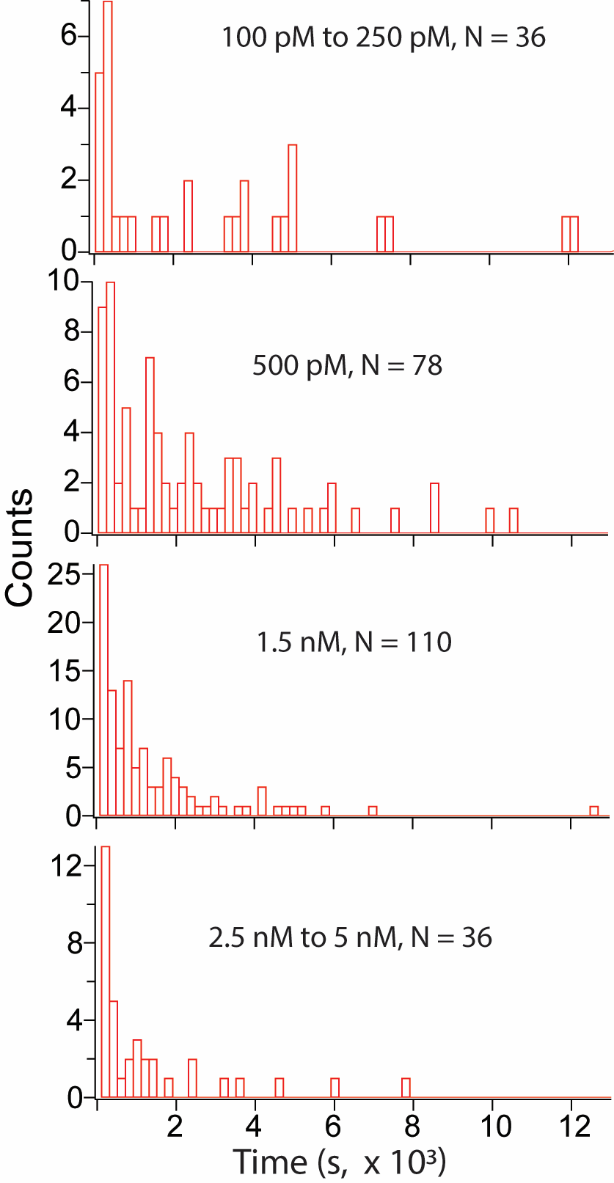

**Figure S3.** Distributions of time intervals between two measurable consecutive events.

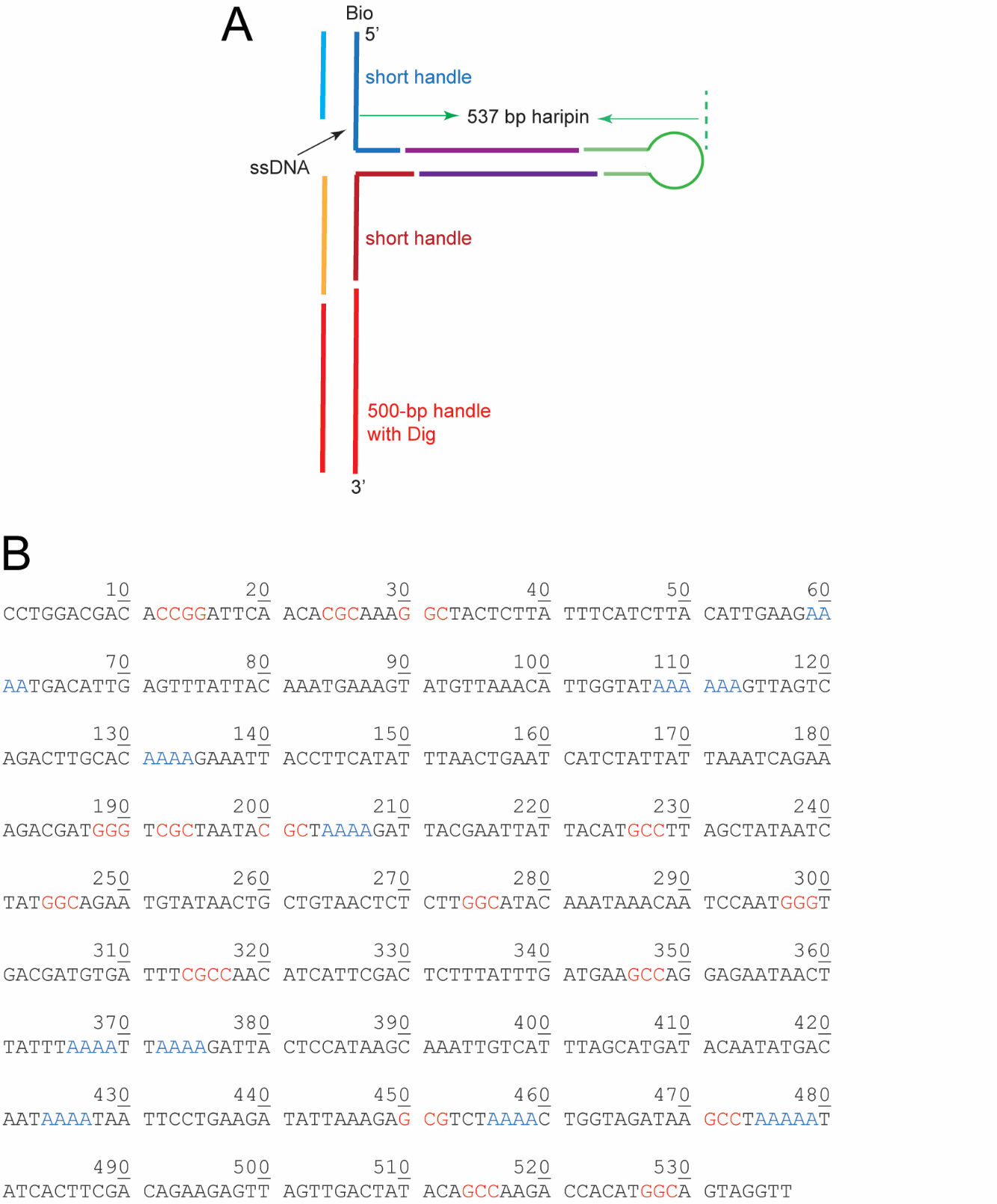

**Figure S4.** The 537-bp DNA hairpin construct. A) Schematic drawing of the construct. B) Sequence of the 5’ half of the unfolded 537-bp hairpin. Stretches with three or more consecutive GC pairs highlighted in red and poly-A stretches with four or more A in blue.

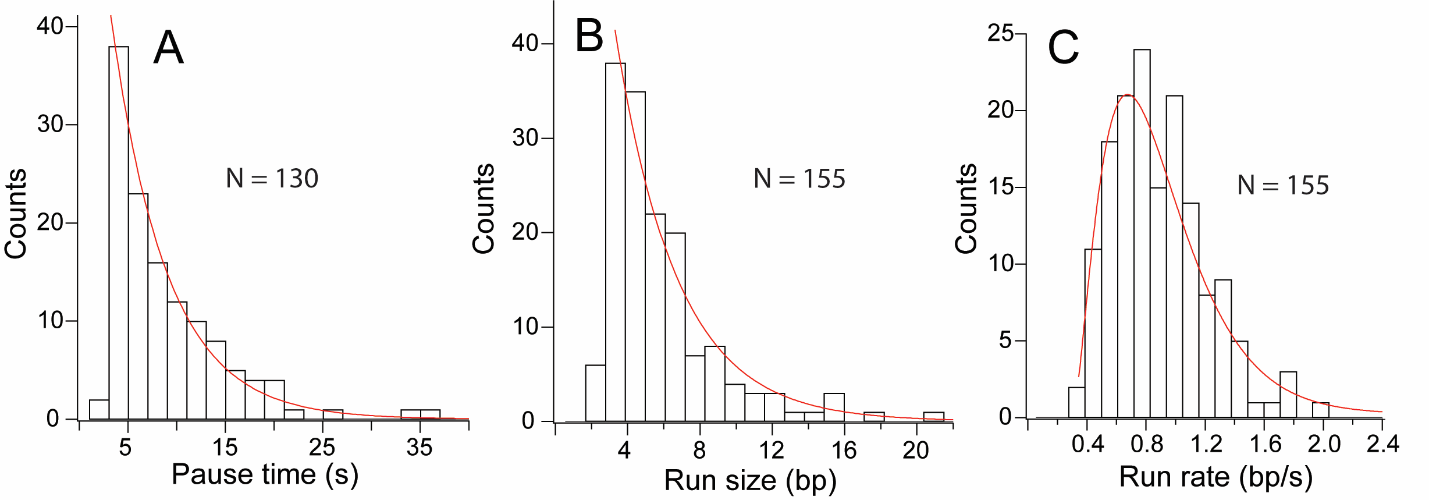

**Figure S5.** Analysis of individual translocation with refolding events. A) Histogram of pause times fitted with exponential curve. B) Histogram of run sizes fitted with exponential curve. C) Histogram of run rates fitted with gamma distribution fit.

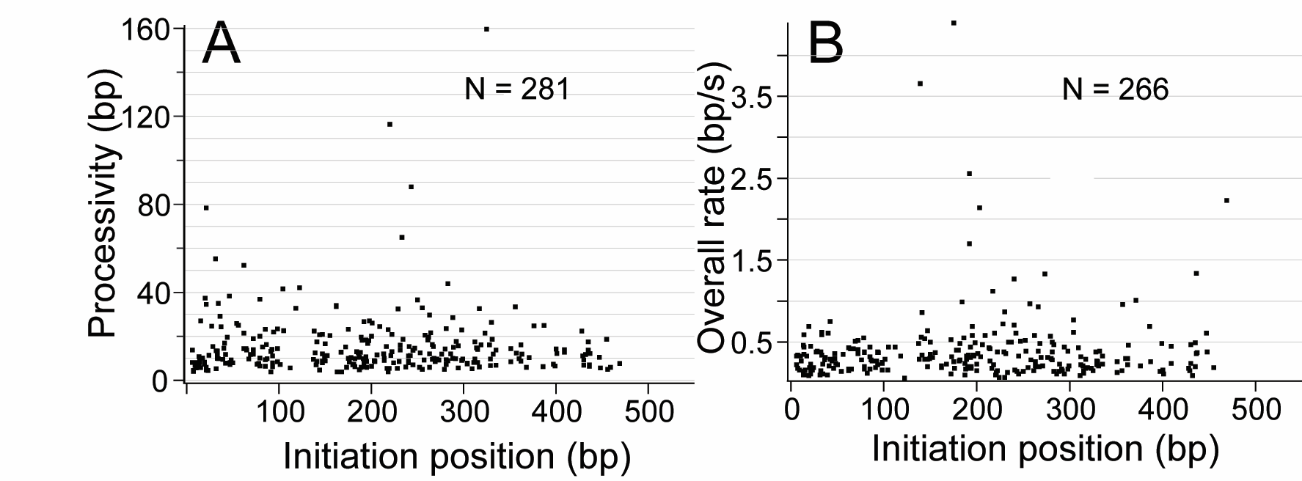

**Figure S6.** Processivity (A) and overall rate (B) of middle unwinding vs. initiation positions.
